# Non-rapid eye movement sleep determines resilience to social stress

**DOI:** 10.1101/2022.06.01.494280

**Authors:** Brittany J. Bush, Caroline Donnay, Eva-Jeneé A. Andrews, Darielle Lewis-Sanders, Cloe L. Gray, Zhimei Qiao, Allison J. Brager, Hadiya Johnson, Hamadi C.S Brewer, Sahil Sood, Talib Saafir, Morris Benveniste, Ketema N. Paul, J. Christopher Ehlen

**Affiliations:** Neuroscience Institute, Morehouse School of Medicine; Atlanta, Georgia, United States; Behavioral Biology Branch, Center for Military Psychiatry and Neuroscience, Walter Reed Army Institute of Research; Silver Spring, Maryland, United States; Department of Integrative Biology and Physiology, University of California; Los Angeles, California, United States

## Abstract

Resilience, the ability to overcome stressful conditions, is found in most mammals and varies significantly among individuals. A lack of resilience can lead to the development of neuropsychiatric and sleep disorders, often within the same individual. Despite extensive research into the brain mechanisms causing maladaptive behavioral-responses to stress, it is not clear why some individuals exhibit resilience. To examine if sleep has a determinative role in maladaptive behavioral-response to social stress, we investigated individual variations in resilience using a social-defeat model for male mice. Our results reveal a direct, causal relationship between sleep amount and resilience—demonstrating that sleep increases after social-defeat stress only occur in resilient mice. Further, we found that within the prefrontal cortex, a regulator of maladaptive responses to stress, pre-existing differences in sleep regulation predict resilience. Overall, these results demonstrate that increased NREM sleep, mediated cortically, is an active response to social-defeat stress that is both necessary and sufficient for promoting resilience. They also show that differences in resilience are strongly correlated with inter-individual variability in sleep regulation.

## Introduction

The links between sleep, neuropsychiatric illness, and responses to stress have been extensively documented, but are poorly understood. Sleep disorders are a debilitating feature of neuropsychiatric conditions (for a review see *1*), and social stress is known to cause or exacerbate both sleep and neuropsychiatric disorders (*2-5*). Notably, both self-reported sleep disturbances and objective polysomnographic recordings are predictive of the development of stress-induced disorders such as post-traumatic stress disorder, depression, and anxiety (*6-11*). In some cases, both humans and animals show resilience to the adverse effects of stress. The degree of resilience is highly variable between individuals for reasons that are not entirely clear *(12, 13)*. The experiments reported here test two hypotheses: 1) that sleep amount plays a determinative role in the response of an individual to stress; and 2) that inter-individual variability in sleep response to stress result in individual differences in resilience.

To investigate these hypotheses, we used a well-established model of social conflict in mice wherein rodents are socially stressed in the home cage of a larger conspecific. This model has been extensively used to model core aspects of human pathologies with high face, construct, and etiological validity (*14-18*). An important aspect of this social-defeat model is that not all socially-stressed animals respond equally well—some mice are susceptible and others are resilient to the effects of stress (*17*). To determine the role of sleep in resilience to social-defeat stress, we have experimentally altered sleep amount; to examine the role of inter-individual variability in sleep, we have conducted detailed examinations of sleep and sleep-response to social-defeat stress in both resilient and susceptible mice.

## Results

We first looked to determine if sleep was necessary for resilience to social-defeat stress. One cohort of mice received daily bouts of social stress at the onset of darkness. In these mice, social interaction (measured by interaction ratio, see methods for details) was assessed one day before and one day after social-defeat stress (**Fig. 1A**). As expected (*17*), half of these mice were resilient (interaction ratio > 1.1) to the effects of social defeat stress (4 of 8 mice were resilient; **Fig. 1B, C**; two susceptible—interaction ratio <1, two undefined—interaction ratio between 1.1 and 0.9). A second cohort of mice was sleep restricted (8h) at light onset during the ten consecutive-days of social-defeat stress (**Fig. 1A**). In contrast to the mice without sleep restriction, no sleep-restricted mouse was resilient; furthermore, all sleep-restricted mice showed decreased social interaction after social-defeat stress (**Fig. 1B, C;** five susceptible, two undefined – interaction ratio between 1.1 and 0.9). Differences in social interaction were not due to changes in ambulatory activity, as all treatment-groups showed similar levels of activity during the social-interaction testing (**Fig. 1E**). In a separate cohort of sleep-deprived mice we assessed fecal corticosterone to estimate stress levels (**Fig. 1D**). These findings show that hypothalamic-pituitary adrenal axis stress-pathways are not significantly activated by sleep restriction (**Fig. 1B**). Collectively, these results demonstrate that sleep restriction increases susceptibility to social-defeat stress and provides evidence that sleep is necessary for resilience to social-defeat stress.

**Figure 1.**
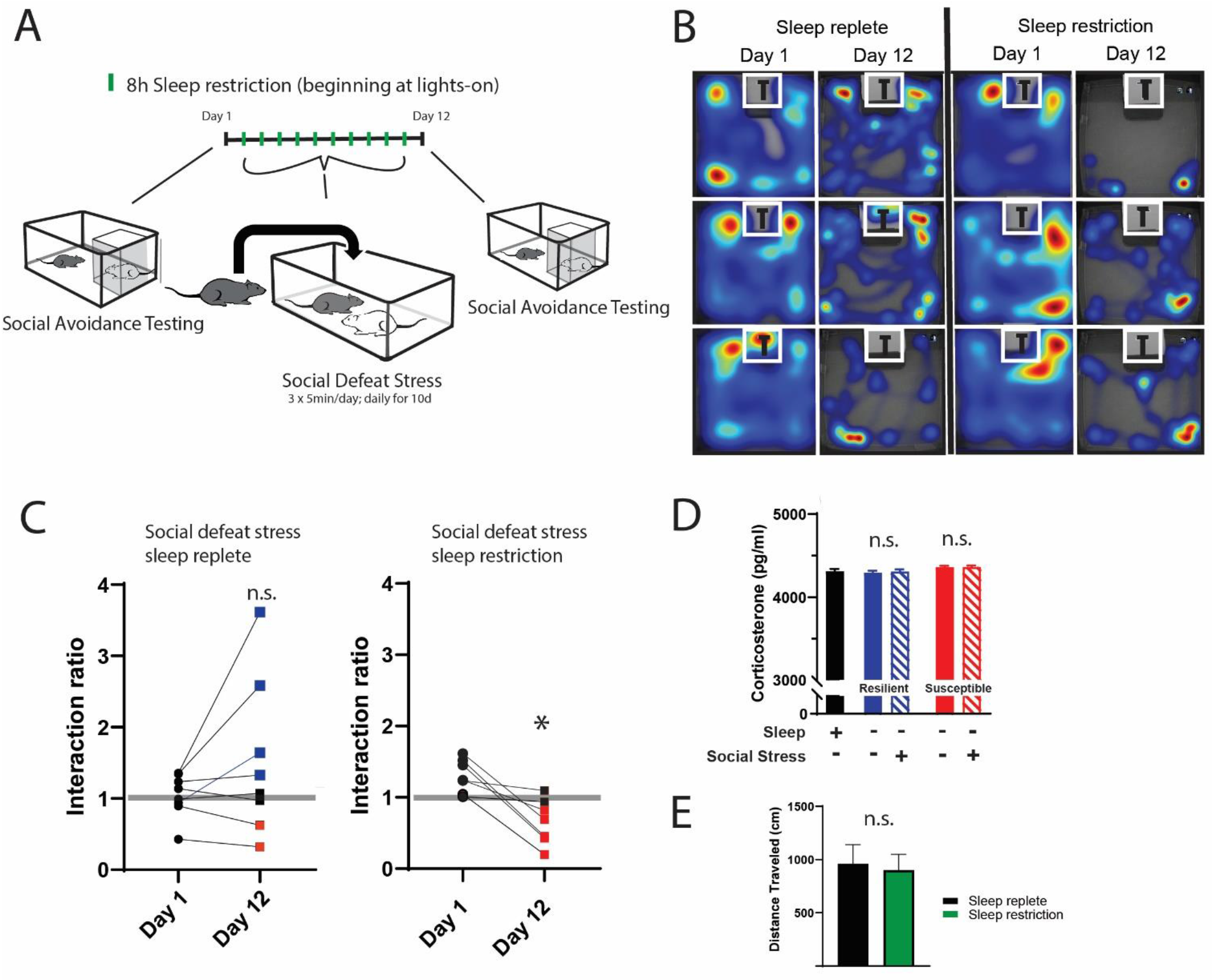
Daily sleep-restriction prevents resilience to social defeat stress. One cohort of mice received sleep restriction (8-h, beginning at light onset) on each day of social defeat stress; a second cohort (sleep replete) received only social defeat stress (A; 10 days total, sleep restriction procedure outlined in Methods). As expected in the sleep replete cohort, roughly equal amounts of resilience and susceptibility occurred after social-defeat stress (B, C). In contrast, no mouse that underwent sleep restriction was resilient to social-defeat stress (B, C). Neither the stress response (D; indicated by fecal corticosterone) nor the distance moved during behavioral testing (E) was significantly altered by sleep deprivation. (B) heatmaps showing the time and location of representative mice during three-minute social-avoidance test both before and after ten days of social-defeat stress. (C) interaction ratios calculated from the heatmaps in B; black circles—pre-stress, red box—susceptible, blue box—resilient, black box—undefined; social avoidance was expressed as an interaction ratio based on the time (t) spent interacting (near white box) with a caged, novel CD1 target mouse vs. an empty cage (interaction ratio = *t*_*e*_ / *t*_*f*_); sleep replete—Student’s paired *t*, t(7)=1.54, *p* = 0.17; sleep restriction—Student’s paired *t, t*(6)=4.02, *p* = 0.007; sleep replete—n = 8, sleep restriction—n = 7. (D) ANOVA, F(4,19)=1.12, p = 0.37; n = 8. (E) Student’s t, t(14)=0.74, p = 0.47. Data points represent mean ± SEM.

To further investigate the role of sleep in resilience, we increased sleep amount using a novel method that avoids the use of somnogenic drugs (**Fig. 2A**). Because sleep is initiated by GABAergic cells projecting from the preoptic area (POA), we used a designer-receptor exclusively activated by designer-drugs (DREADD) to activate these POA neurons. Four weeks after delivering DREADD to the POA by adeno-associated virus, clozapine-n-oxide (CNO) was used to activate the DREADD and increase sleep. This was validated in a separate cohort of mice at zeitgeber time 10 (ZT, ZT 12 = lights-off, **Fig. 2C**), where a single injection of CNO increased non-rapid eye movement (NREM) sleep for approximately 6-hours. Single, daily injections (administered ZT 1–2) of CNO across the 10-day social-defeat paradigm enhanced total sleep by 79.3 ± 12.5 (mean ± SEM) min per day over control mice, (Fig. **2B**) including 114.6 ± 14.2 minutes per day of NREM sleep. Part of this NREM increase occurred at the expense of rapid eye movement (REM) sleep; REM sleep was reduced by 31.6 ± 4.7 min per day. Mice expressing the DREADD (increased sleep) had significantly increased interaction ratios (increased resilience) after social defeat stress, when compared to CNO-treated control mice expressing EGFP (**Fig. 2E**). Notably, all DREADD-expressing mice (increased sleep) were resilient (interaction ratio > 1.1), whereas only half of control-treated mice were resilient. These results provide evidence that increased sleep confers resilience and has a protective influence during exposure to social-defeat stress; supporting our hypothesis that sleep is both necessary and sufficient for resilience to social-defeat stress.

**Figure 2.**
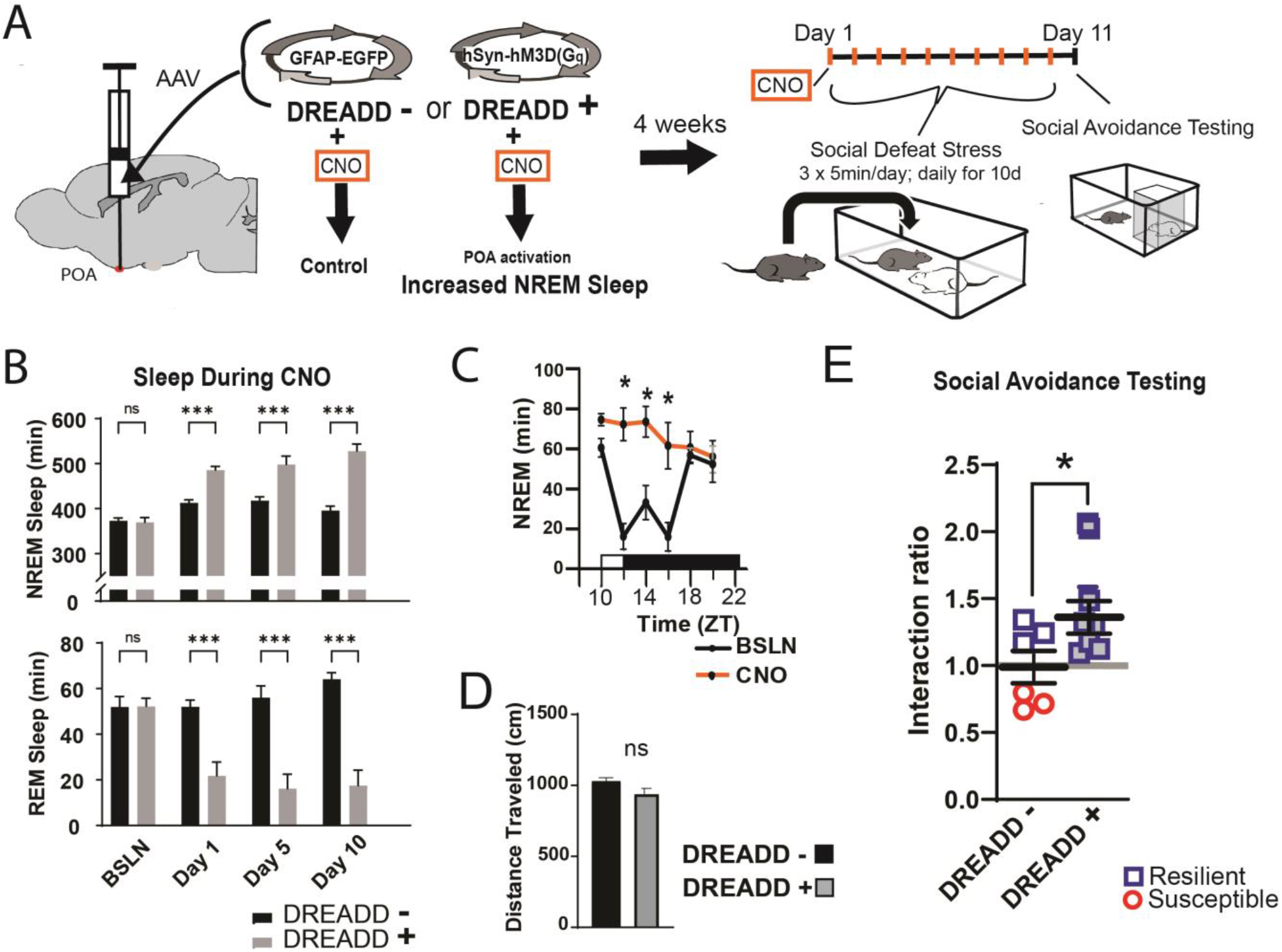
Increased sleep promotes resilience to social defeat stress. Adeno-associated viral vectors (AAV2) encoding either an excitatory (G_q_) designer receptor exclusively activated by designer drugs (DREADD), or enhanced green fluorescent protein (EGFP) as a control, were delivered to the preoptic area (POA) by intracranial microinjections. After four weeks, i.p. injections of clozapine N-oxide (CNO) were used to activate receptors expressed in the POA (A). In a validation study, chemogenetic activation of the POA significantly increased NREM sleep for six hours (compared to undisturbed sleep on the previous day) following a single injection of the agonist CNO at zeitgeber time 10 (C; ZT 10, ZT12 = lights off). A separate cohort of mice expressing DREADD, or EGFP control, was exposed to 10 days of social defeat stress (ZT12–13) with single, daily, i.p. injection of CNO at lights on (ZT 1–2; A). NREM sleep was significantly increased by daily injections of CNO in mice expressing the excitatory DREADD, but not in mice expressing the control DREADD (B). No mouse expressing the excitatory DREADD was susceptible to the effects of social defeat stress (E). Mice expressing the control construct displayed both susceptible and resilient behavior as expected (E). The distance moved during behavioral testing was not significantly affected by POA activation (D). (B) repeated measures ANOVA: NREM main effect of CNO—F(1, 35) = 83.37, *p* < 0.0001; interaction effect—F(3, 35) = 13.43, *p* < 0.0001; REM main effect of CNO—F(1, 29) = 72.92, *p* < 0.0001; interaction effect—F(3, 29) = 10.5, *p* < 0.0001; *, *p* ≤ 0.001, Holms Sidak’s multiple comparison; n = 6, DREADD-; n= 9 DREADD +. (C) repeated measures ANOVA: main effect of CNO—F(1,8) = 12.82, *p* = 0.008; interaction effect—F(5, 40) = 8.04, *p* < 0.0001; *, *p* ≤ 0.001, Holms Sidak’s multiple comparison. (D) Student’s t, t(11)=1.82, p = 0.087. (E) Student’s t, t(11)=2.157, p = 0.027. Data points represent mean ± SEM; * = *p* 0.05.

We next looked for the differences in sleep amount and architecture between resilient and susceptible populations. Analysis of sleep before and after social defeat stress revealed dramatic changes in post-defeat sleep, but only in resilient mice (**Fig. 3A**,**B & C**). Both the REM and NREM sleep of resilient mice were increased throughout the active/dark period following defeat, when compared to baseline sleep (**Fig. 3C**). These sleep changes after defeat included an increase in total sleep amount over the 24-hour period (**Fig. 3D**) and a reorganization of sleep from typical baseline patterns. Indeed, most of the post-defeat changes in total-sleep occurred during the active/dark period (**Fig. 2D**). Control mice exposed to novel cages over 10-days did not have this reorganization of REM and NREM sleep. Instead, control mice had a modest increase in NREM and REM sleep during the dark period (**Fig. S1**), showing that the effect of a novel environments on sleep was minor. NREM slow-wave activity (power 0.5–4Hz) and theta activity (6–10Hz), were also altered in resilient mice (**Fig. 3E**). NREM slow-wave activity is a standard measure of sleep-intensity. These differences in SWA reveal underlying differences in sleep regulation between resilient and susceptible populations of mice. Collectively, our data demonstrate that sleep is both increased and reorganized after social defeat stress exclusively in resilient mice and suggest that differences in sleep regulation may underlie these sleep changes.

**Figure 3.**
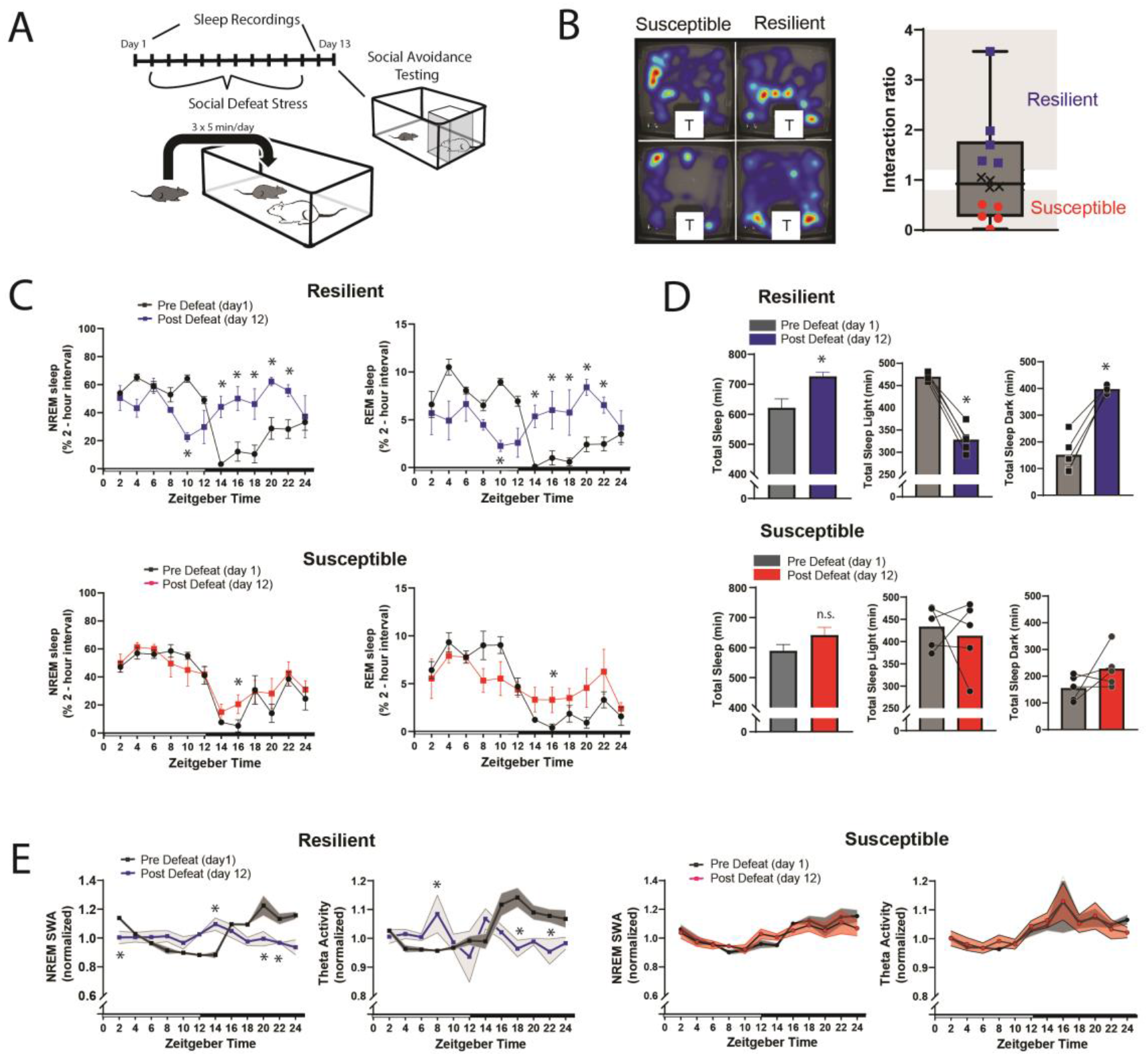
Sleep is reorganized only in mice resilient to social defeat stress. 24-hour sleep recordings were performed both before and following 10-days of social defeat stress (A). Sleep in animals identified as resilient (B) was significantly reorganized (C) and increased (D) following social defeat stress. This change in resilient mice included a significant decrease in total sleep during the light period and increased total sleep during the dark period (D). Animals identified as susceptible (B) showed little change in sleep architecture or amount (C, D). Sleep changes in resilient animals included a flattening of the normal curve in sleep intensity (E; NREM slow-wave activity, SWA: power density 0.5–4 Hz). This change was accompanied by changes in higher frequencies during NREM sleep, theta activity was increased during the day and decreased during the night (E). In contrast, no significant change in either NREM slow-wave or theta activity was observed in susceptible animals (E). (B) left, representative heatmaps of social avoidance testing; warmer colors indicate increased time; T=caged mouse; right, interaction ratios, X indicates interaction ratios between 0.9 and 1.1 that were excluded from sleep analysis. (C) resilient, repeated measures ANOVA: NREM: main effect of time—*F*(11, 88) = 4.86, *p* = 0.0001; main effect of day—*F*(1,8) = 11.71, p= 0.009; interaction—*F*(11,88) = 8.02, p = 0.0001. REM: main effect of time—*F*(11, 88) = 3.17, *p* = 0.0012; main effect of day—*F*(1,8) = 0.95, p=0.358; interaction—*F*(11,88) = 6.7, p = 0.0001. Susceptible, repeated measures ANOVA: NREM main effect of time—*F*(11, 88) = 8.581, *p*<0.0001; main effect of day—*F*(1, 8) = 2.925, *p* = 0.1256; interaction—*F*(11, 88) = 1.429, *p* = 0.1742; n = 12. (D) resilient, Student’s paired t: total sleep—*t*(5) = 5.09, *p* = 0.007; light—*t*(5) = 14.62, *p* = 0.0001; dark—*t*(5) = 8.15, *p* = 0.0012. (E) resilient, SWA: main effect of time—F(11, 84) = 4.482, p < 0.0001; main effect of susceptibility—F(1, 8) = 2.84, p = 0.14; interaction—F (11, 84) = 6.47, p = 0.0001. NREM theta activity: main effect of time—F(11, 89) = 1.97, p = 0.042; main effect of susceptibility—F(1, 89) = 5.04, p = 0.027; interaction—F (11, 89) = 4.39, p<0.0001. susceptible: main effect of time— F(11, 84) = 6.31, p < 0.0001; main effect of susceptibility—F(1, 8) = 0.64, p = 0.45; interaction—F (11, 83) = 1.31, p = 0.23; *, *p* ≤ 0.05, Holms Sidak’s multiple comparison. Data points represent mean ± SEM except for B which presents median, 25th to 75th percentiles and min/max values.

We also investigated whether sleep-regulatory differences underlie the sleep responses of resilient and susceptible mice. Sleep is homeostatically regulated, as NREM sleep amount and intensity (slow-wave activity, SWA) are proportional to the duration of prior wakefulness (*19, 20*). A standard method for investigating this homeostatic process involves restricting sleep and then measuring the resulting changes in NREM amount and intensity. To investigate sleep-regulatory differences that may underlie resilience to social-defeat stress, we used a sleep restriction paradigm both before and after exposure to social-defeat stress (**Fig. 4A**). Prior to any stress exposure, mice later identified as susceptible showed increased sleep-recovery (**Fig. 4B; 4C, top row**), and increased NREM SWA **(Fig. 3 G)**, after six-hours of sleep restriction; mice later identified as resilient did not show these changes. This difference was not caused by the amount of sleep lost, as both behavioral phenotypes lost similar amounts of NREM sleep during sleep restriction (**Fig. 4C**, bottom row). In addition, the increased recovery-response in susceptible mice remained when NREM sleep was normalized for each mouse (to the amount of sleep lost over six-hours and 18-hours of recovery; **Fig. 4C**, bottom row). After exposure to social stress, the overall patterns in NREM-sleep and NREM SWA were similar to pre-stress values (**Fig. 4D, F bottom row**). Notably, NREM SWA in susceptible mice was greater than resilient mice, both before (i.e. under baseline conditions) and after exposure to social-defeat stress (**Fig. 4 E & F, top row**). Both behavioral phenotypes showed equivalent increases in REM-sleep and total sleep (**Fig. 4D, top row**). These findings demonstrate that sleep and sleep-regulatory changes in resilient mice are not simply caused by social-defeat stress, instead, differences in sleep regulation exist prior to social-defeat stress and predict resilience. Collectively, our findings suggest that pre-existing differences in sleep regulation determine the sleep-response to social-defeat stress; these sleep responses, in turn, determine resilience.

**Figure 4.**
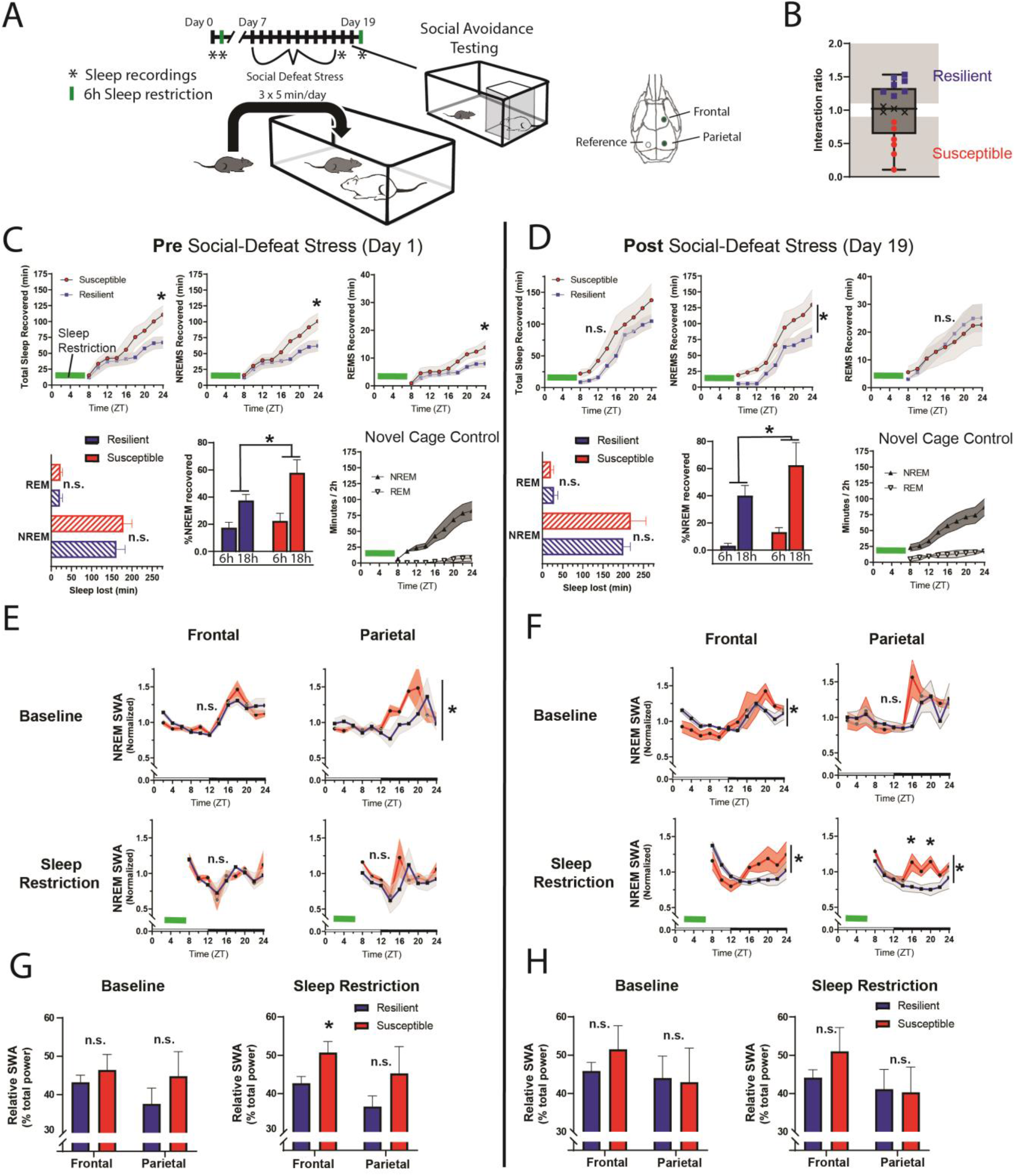
Differences in sleep regulation, prior to stress exposure, predicts resilience to social defeat stress. Sleep-regulatory mechanisms were assessed with a six-hour sleep restriction performed both before and after 10-days of social defeat stress (A). Prior to any stress exposure, mice later identified as susceptible showed increased sleep-recovery to six-hours of sleep restriction (B; C, top row) when compared to mice identified as resilient (B; C, top row). This pattern was present for both NREM and REM sleep (C, top row). Susceptible and resilient mice lost similar amounts of NREM and REM sleep during sleep-restriction (C, bottom row) and the increased recovery response for susceptible mice persisted when NREM sleep recovered was normalized to the amount of sleep lost (C, bottom). After social defeat stress, susceptible animals continued to show increased NREM-sleep recovery, but not REM-sleep recovery (D, top row). Sleep intensity (NREM slow-wave activity, SWA) was higher in susceptible animals prior to social defeat stress during both baseline and following sleep restriction (E, G). This increased sleep intensity in susceptible mice persisted and was more prominent after social defeat stress (F, H). (B) X indicates interaction ratios between 0.9 and 1.1 that were excluded from sleep analysis. (C) Total sleep: repeated measures ANOVA main effect of time (MET)—F(8, 80) = 53.78, p<0.0001; main effect of susceptibility (MES)—F(1, 10) = 2.05, p = 0.18; interaction (IST)—F (8, 80) = 5.14, p<0.0001. NREM sleep, MET—F(8, 80) = 50.75, p < 0.0001; MES—F(1, 10) = 1.92, p=0.19; IST—F (8, 80) = 4.77, p < 0.0001. REM sleep, MET—F(8, 80) = 36.88, p < 0.0001; MES—F(1, 10) = 2.73, p = 0.13; IST—F (8, 80) = 3.32, p<0.0025. Sleep lost, Student’s paired t, NREM—*t*(7) = 0.99, *p* = 0.36; REM—*t*(7) = 0.92, *p* = 0.37. % recovered: MET—F(8, 80) = 41.45, p<0.0001; MES—F(1, 10) = 1.58, p=0.24; IST—F (8, 80) = 3.27, p=0.0029. (D) NREM sleep: MET—F(8, 56) = 49.2, p<0.0001; MES—F(1, 7) = 3.96, *p* = 0.049; IST—F (8, 56) = 1.86, p=0.23. REM sleep: MET—F(8, 56) = 24.58, p<0.0001; MES—F(1, 7) = 0.02, p = 0.87; IST—F (8, 56) = 0.48, p=0.86. (E) Baseline, parietal: MES—F(1, 95) = 4.21, *p* = 0.043. (F) Frontal baseline: IST—F(11, 75) = 2.13, *p*=0.028; frontal sleep restriction, IST—F(8, 63) = 2.13, *p* = 0.028; parietal sleep restriction, IST—F(8, 63) = 2.84, p = 0.009). * = *p*≤0.05, Holms Sidak’s multiple comparison; n = 13. Data points represent mean ± SEM with the exception of panel B which presents median, 25th to 75th percentiles and min/max values.

The ventromedial prefrontal cortex (vmPFC; prelimbic, PrL and infralimbic, IL cortex) is important in regulating stress resilience (*21*) and is also sensitive to the effects of sleep restriction (*22, 23*). The technique used for our EEG recordings (epidural screw electrodes) does not provide optimal spatial resolution. To better investigate local differences in sleep, we used local field potential recordings (LFP) from electrodes deep in the vmPFC. When compared to susceptible mice, mice later identified as resilient, showed a significant increase in baseline LFP power density in the slow-wave range (≤ 5 Hz) before social defeat stress (**Fig. 5A**). This SWA in baseline NREM sleep was found at both major vmPFC subregions targeted by our LFP-electrodes. The difference was most prominent in the PrL cortex and also detected in the IL cortex. Notably, no power differences were found in our EEG recordings (although a trend may be present ≤ 2Hz; **Fig. 5A, S2**). These findings suggest that local differences in NREM slow-wave activity, between susceptible and resilient mice, exist in the vmPFC before exposure to social stress and can predict resilience.

**Figure 5.**
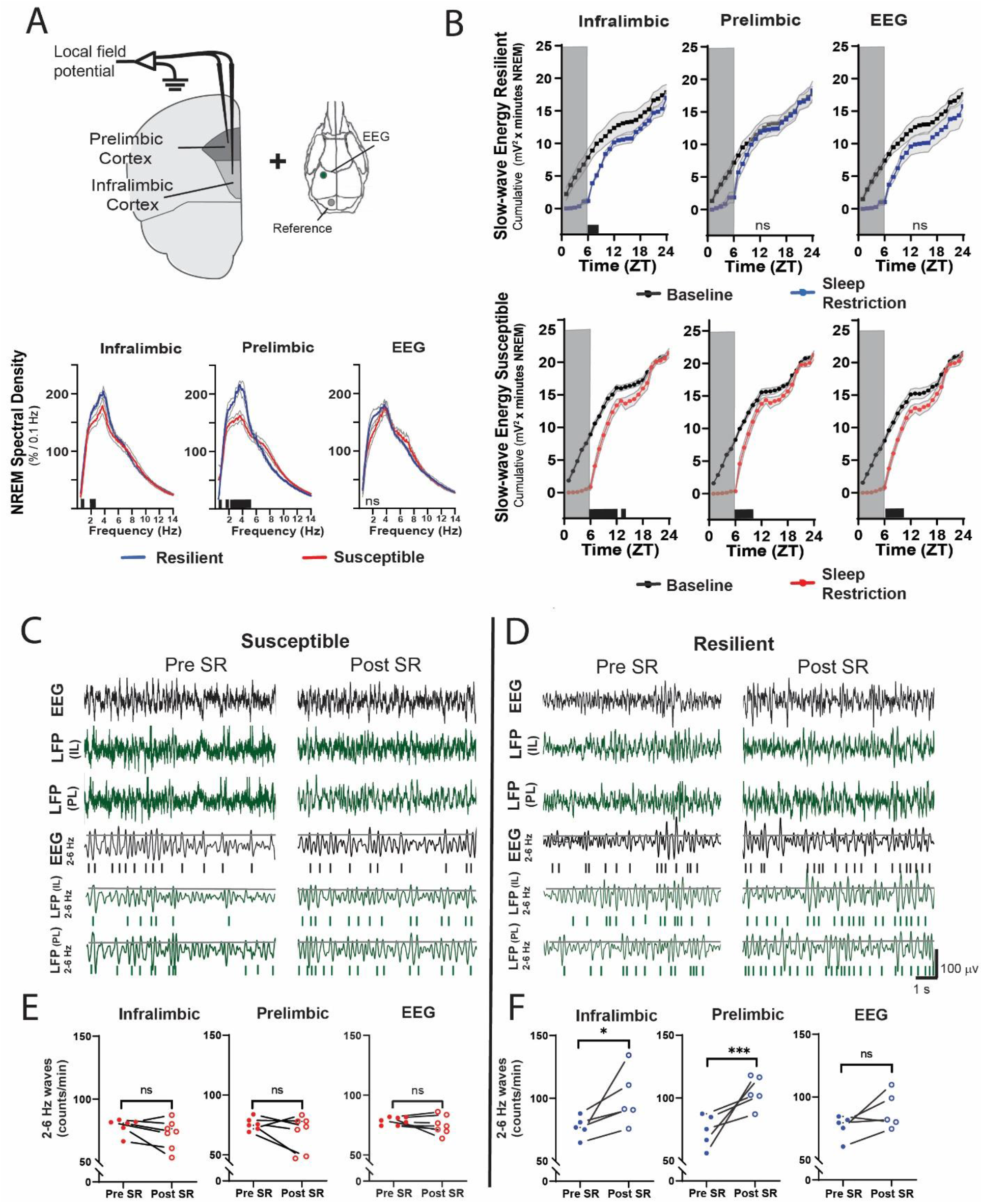
Sleep changes in the ventromedial prefrontal cortex (vmPFC) predict resilience to social defeat stress. Local field potential (LFP) in the vmPFC and epidural electroencephalographic recordings (EEG; A, top row; black bars on x-axis = p≤0.05) were simultaneously obtained from mice before ten-days of social defeat stress. 24-hour LFP/EEG recordings were immediately followed by six-hour sleep restriction as in Figure 4A (no post-defeat restriction). NREM sleep intensity (power density > 4 Hz) in both the prelimbic and infralimbic LFP were significantly higher in mice later identified as resilient (vs. susceptible mice; A, bottom row); notably, these differences in sleep intensity were not evident in the EEG (A, bottom row). After six-hours of sleep restriction resilient animals recovered at a faster rate than susceptible animals. This recovery is observed in cumulative NREM slow-wave energy (delta band = 0.5-4 Hz; 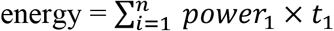) and resilient mice took significantly less time to reach baseline levels (B). The incidence of 2–6 Hz waves in the waking EEG, a marker of sleep-pressure during waking, significantly increased in sleep restricted mice identified as resilient (D). This occurred in both LFP and EEG recordings and indicates a normal accumulation in sleep-pressure. Mice later identified as susceptible (C) did not show this increased wave-incidence, thus, indicating a lack of sleep-pressure accumulation. (A) shaded areas are SEM; grey boxes indicate sleep restriction; ANOVA; prelimbic interaction—*F* (294, 2950) = 4.55, *p* < 0.0001; infralimbic main effect of susceptibility—F (294, 2950) = 4.22, p < 0.0001; EEG, interaction effect—*F* (144, 1450) = 0.9175, p = 0.74; Holms Sidak’s multiple comparison; n = 12. (B) repeated measures ANOVA interaction effect—susceptible mice: infralimbic LFP F(17, 187) = 11.13, *p* < 0.0001; prelimbic LFP F(17, 187) = 5.71, *p* < 0.0001; EEG F(17, 187) = 6.75, *p* < 0.0001; resilient mice: infralimbic LFP F(17, 136) = 3.71, *p* < 0.0001; black bars on x-axis = *p*<0.05, Holms Sidak’s multiple comparison. (C, D) top rows— representative, raw EEG (black) and LFP (green) recordings; middle rows—filtered EEG (black) and LFP (green) signals (2–6 Hz) with threshold for counting identified (70^th^ percentile, black line, see methods for details). (E, F) Lower plots—average wave-incidence for the 1-hour period immediately before and immediately after sleep restriction; Student’s paired t, *, *p* = .03; ***, *p* = .0008.

Next, we looked at recovery responses to sleep restriction before exposure to social defeat stress. Slow-wave energy was recovered in significantly less time for resilient animals, indicating a more efficient sleep-regulatory response (**Fig. 5B**). Slow-wave energy returned to baseline levels immediately after sleep restriction in the PrL LFP and EEG of resilient mice, and within two-hours in the IL LFP. Susceptible mice, in comparison, were not fully recovered for up to 8-hours (**Fig. 5B**). We also examined the LFP during wake, both before and after sleep restriction. Low-frequency waveforms (2–6 Hz) increase in number and amplitude with sleep-pressure accumulation (*24, 25*). These waveforms are thought to represent local sleep-like events encroaching into wakefulness (i.e. sleepiness). Counting the occurrence of the largest amplitude 2–6Hz waveforms (I_2-6_) revealed that I_2–6_, and thus waking sleep-pressure, was increased in resilient mice after sleep-restriction. Susceptible mice showed no significant change in I_2–6_ within the awake LFP (**Fig. 5C**). To further assess local differences in the vmPFC, we calculated phase coherence between each LFP electrode and the global EEG. We found phase coherence below 8 Hz increased in resilient mice during all conditions including: baseline, recovery from sleep restriction and after social defeat stress; however, this difference was only found in the IL cortex (**Fig. 6**). Importantly, the 24-hour average of these coherence values, in slow-wave frequencies, was significantly and positively correlated with social interaction ratios for the IL cortex, but not for the PrL cortex (**Fig. 6B**). These LFP studies further support our conclusion that local differences in sleep-regulatory processes within the vmPFC exist before stress and predict resilience.

**Figure 6.**
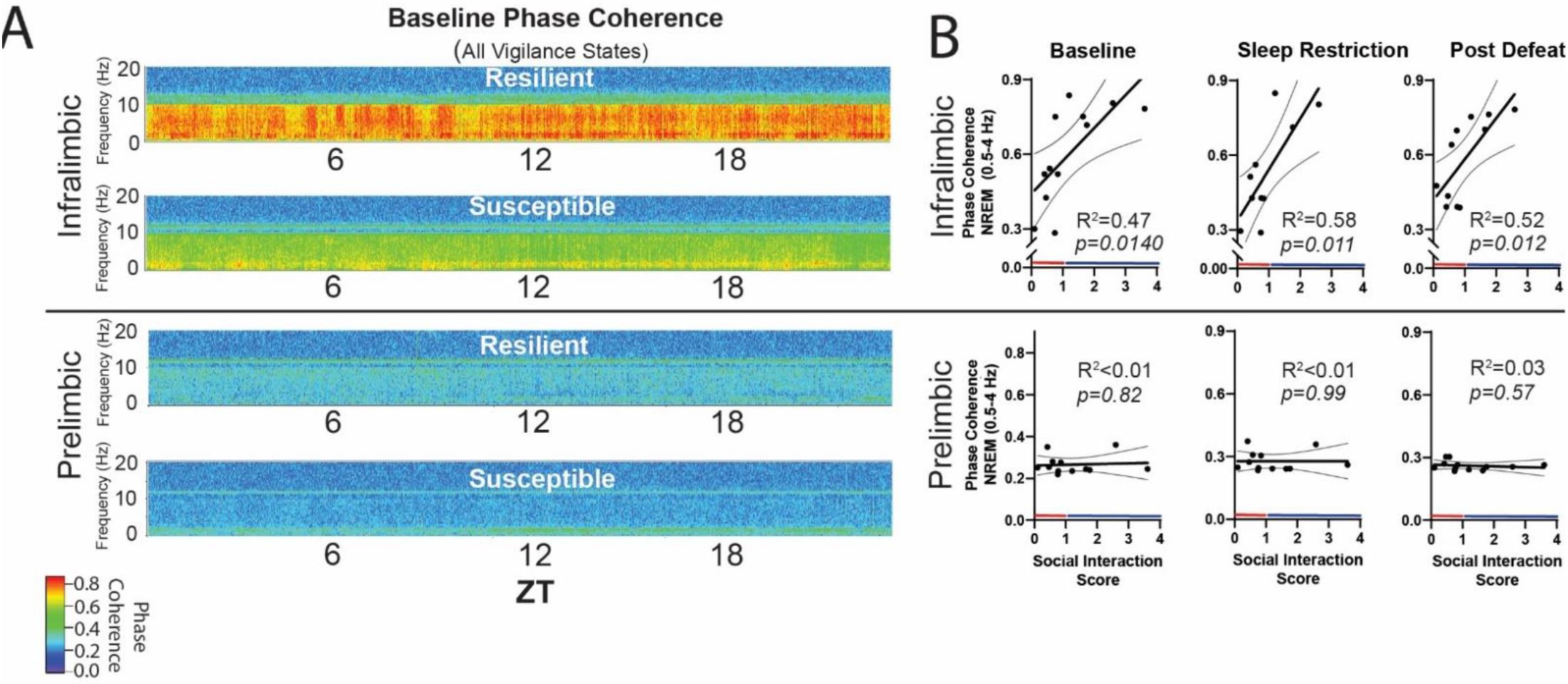
NREM coherence in the 0.5–4Hz range predicts resilience to social defeat stress. Local field potential (LFP) in the ventromedial prefrontal cortex and epidural electroencephalographic recordings (EEG) were simultaneously obtained from mice before and after ten-days of social defeat stress (see Fig. 5). Coherence below 8 Hz between the infralimbic (IL) LFP and EEG was significantly increased across the 24-hour day in undisturbed, resilient mice (compared to susceptible; A, top; warmer colors represent increased coherence). This increased coherence with the EEG below 10 Hz was also visible in the prelimbic (PrL) LFP, but this effect was reduced and not significant (A, bottom; compared to susceptible). A similar pattern of increased coherence in resilient animals was observed after six-hours of sleep restriction and after 10-days of social defeat stress (data not shown). Notably, IL-coherence (averaged over 24-hours) predicted social interaction ratios (B, top) both before (B, left) and after (B, middle) social defeat stress and during recovery from six-hours of sleep restriction (B). These correlations were not observed for the PrL cortex (B, bottom). (A) heatmaps of coherence over 0.1 Hz intervals in 10-minute bins; IL, repeated measures ANOVA: main effect of susceptibility—F(1, 10) = 19.81, *p* = 0.0024; PrL, main effect of susceptibility—F(1, 10) = 0.188, p = 0.09 n.s; n = 12. (B) least-squares regression line with 95% confidence interval and goodness of fit (R^2^); colors represent resilience (blue) or susceptibility (red) based on interaction ratio; non-zero slope of least-squares regression line, baseline—F(1,10) = 8.87, sleep restriction—F(1,10) = 10.99, post social defeat—F(1,10) = 9.83; *p* values provided in plots.

## Discussion

We have applied behavioral, electrophysiological and chemogenetic approaches to investigate a causal link between pre-existing sleep-differences and behavioral responses to stress. We demonstrate through sleep restriction that sleep is required for resilience to social defeat stress; further, our POA-activation findings demonstrate that specifically increasing sleep amount increases resilience. Thus, sleep is both necessary and sufficient for resilience to social stress. Furthermore, sleep recordings obtained before and after social-defeat stress reveal that increases in sleep are exclusive to resilient animals. Together, these data lend strong support for the essential role of sleep in conferring resilience.

Several previous studies have revealed that social stress in both mice and rats causes increased sleep intensity (*2*); notably, this increased sleep intensity also occurs in the ‘winners’ of social conflict (*26*). Meerlo et al., 1997 have suggested that the specific nature of preceding wakefulness as well as the duration of prior wakefulness is important in sleep responses (*2*). Thus, social defeat stress may demand increased recovery sleep because it represents a more intense form of wakefulness (*2*). Indeed, the effect of waking-context on sleep regulation may extend beyond stress; recent findings indicate that repetitive tasks may represent less-intense wakefulness that, in turn, leads to less intense sleep (*26, 27*). In this context, the altered sleep regulation of susceptible mice in our experiments may render them incapable of responding with increased sleep during or after social defeat stress, whereas resilient animals adequately recover from this intense wakefulness. Although the ability of social defeat stress to alter sleep regulation has been reported (*28, 29*), our studies demonstrate that differences in sleep regulation exist before exposure to stress. The recent finding that sleep fragmentation, a potential indicator of altered sleep regulation, predicts stress susceptibility (*30, 31*) is consistent with this hypothesis; fragmentation before stress exposure was also confirmed in our studies (**Fig. S3, B**; longer NREM bout durations in resilient mice). Thus, the sleep changes, and ultimate behavioral outcomes resulting from social defeat stress, are likely the result of pre-existing differences in sleep regulation. In this context, the intense waking experience of social stress, interacting with differences in sleep regulation between susceptible and resilient mice, may allow resilient animals to recover sleep and leave susceptible animals in a stress-vulnerable, perpetually sleep deprived state.

Susceptible mice appear to be sleep deprived in that they show markers of insufficient sleep after 6-hours of sleep restriction. The deprivation was indicated by increased recovery sleep, both before and after social-defeat stress (**Fig. 4 C, D**); as well as a delayed recovery of slow wave energy (Fig. 5B). Insufficient sleep causes both a reduced ability to cope with stress and negative effects on mood (for a review see (*32*)). Functional magnetic resonance imaging studies reveal that sleep deprivation leads to decreased functional connectivity between the ventromedial prefrontal cortex (vmPFC) and amygdala; furthermore, this decreased functional connectivity is associated with decreased mood (*33-35*). This same brain region, the vmPFC, is known to mediate resilience to social defeat stress (*21, 36-39*). We reasoned that the sleep deprived state of susceptible mice would lead to vmPFC disfunction; thus, leading to decreased connectivity and an inability to inhibit the limbic circuits responsible for regulating behavioral responses to social stress. As predicted, we found major differences between the vmPFC of susceptible and resilient mice. Our LFP recordings reveal that baseline slow wave activity and recovery from sleep deprivation are preferentially enhanced in the vmPFC of resilient animals— before exposure to stress (**Fig. 5A, B**). Wave incidence after sleep restriction, a marker of waking sleep pressure, is also significantly increased in resilient mice (**Fig. 5C–F**). Together, the enhanced buildup of sleep pressure and increased homeostatic response led to faster recovery from sleep restriction in the vmPFC of resilient mice, thus supporting the hypothesis that NREM-related changes in vmPFC are associated with resilience.

In the present study, we used activation of the POA as a method to increase sleep. The POA has a well-established role in the promotion of sleep and multiple cell groups and neuronal subtypes within this region are involved in initiating sleep (*40-42*). Other physiological responses are also regulated by the POA including exploratory and sexual behavior, shivering thermogenesis and body temperature (*43-45*). A recent chemogenetic study activated galanin-expressing neurons in the POA and reported findings similar to those reported here (increased NREM and decreased REM sleep); also reported was a decrease in body temperature (*45*). We did not observe behavioral effects other than sleep and our method was different in that it activated all neurons in the region; however, we cannot completely rule out the occurrence of non-specific effects. We also conducted a detailed histological examination of this region to verify DREADD expression. All mice in the study expressed DREADD in regions of the POA known to initiate sleep (**Fig. S4**) and all responded with increased NREM sleep **(Fig. 2**). Nevertheless, when considered in the context of our other results, especially our finding that sleep restriction decreases resilience (i.e. sleep restriction has the opposite effect of POA activation), our studies provide strong evidence that changes in sleep mediate the changes we observed in resilience.

Sleep regulation in the cortex is known to occur at a local level (*25, 46, 47*), areas that are more active during awake periods are known to have increased sleep intensity during subsequent sleep episodes(*48, 49*).To find if local sleep-differences contribute to resilience, we examined sleep-related changes in two major subdivisions of the vmPFC—the infralimbic (IL) and prelimbic (PrL) cortex. Phase-coherence in the IL cortex (0.5–4 Hz, with the epidural EEG) was significantly and positively correlated with social-interaction ratio (**Fig. 6**). This positive correlation increased after sleep restriction and persisted after social stress. No such positive correlation was found in the PrL cortex. Thus, enhanced phase-locking (i.e. coherence) of the IL cortex with the global EEG (in NREM frequency ranges) predicts resilience. Other subregional differences included enhanced SWA (5A, 0.5–4Hz range) and slow-wave-energy recovery (Fig. **5B**) in the PrL cortex. Collectively, these findings suggest that the sleep-regulatory differences predicting resilience can be further localized to specific subregions of the vmPFC. Both the IL and PrL cortex have a demonstrated importance in resilience to social defeat stress (*21, 36, 50, 51*); thus, the significance of the observed differences between subregions of the vmPFC (e.g. coherence, power density and slow-wave energy) is not yet clear. Nevertheless, the data demonstrate a clear positive correlation between NREM sleep in the vmPFC and resilience.

Our findings indicate a major role for NREM sleep in mediating resilience to social-defeat stress based on several lines of evidence. First, our sleep deprivation paradigm decreased both NREM and REM sleep whereas activation of the POA preferentially increased NREM sleep while REM sleep was reduced. Because REM was decreased in both conditions, with opposite effects on social interaction, REM sleep is unlikely to have large effects on resilience. Furthermore, evidence of altered NREM sleep regulation was most prominent in the vmPFC of resilient mice. In this area, the frequency bands that predominate in NREM sleep were significantly higher and NREM responses to sleep deprivation were significantly enhanced in resilient mice (**Fig. 5 & 6**). Together, these findings strongly implicate NREM sleep in mediating changes in resilience, but do not rule out the involvement of REM sleep.

The effects of stress on behavior, and sleep-responses to stress, vary with sex (*52-54*); thus, it is not possible to predict how our findings relate to females. Female mice are not territorial; therefore, social-defeat stress in females requires alternative defeat-paradigms. These female-defeat paradigms were only recently developed and reported (*54, 55*), which prevented us from considering sex as a biological variable in the present study. Studies in females will be critical to understanding the role of NREM sleep in resilience and are currently underway in our lab.

The present studies show that sleep is an active response to social stress that serves to promote resilience, thus demonstrating a clear causal link between insufficient sleep and maladaptive behaviors. Further, our findings in the vmPFC reveal local changes in sleep that may not be visible in the global EEG. This induced sleep response is dependent on inter-individual variability in sleep regulation and, if sufficient sleep is obtained, likely serves to mitigate the well-established negative effects of sleep loss on CNS function—including cognition and emotion (*56-58*). In addition, the newly demonstrated ability of sleep manipulations to alter behavioral responses to stress offers new possibilities for this mouse model of resilience. This model can be used to investigate the specific mechanisms by which sleep alters stress-induced changes in brain physiology and behavior

## Materials and Methods

### Animals

Male, C57BL/6J mice (Jackson Laboratory, Bar Harbor, ME, USA; 000664) were seven-weeks old at the start of all studies. CD-1 retired male breeders (Charles Rivers, age 3 to 6 months upon arrival) were used as aggressors. All mice were singly housed on shaved, pine bedding upon arrival, maintained on a 12:12 L:D lighting cycle for the remainder of the study and randomly assigned to treatment groups. Food and water were available *ad libitum*. All procedures involving animals received prior approval from the Morehouse School of Medicine Institutional Animal Care and Use Committee (approved protocol 21-02).

### Surgery: EEG and LFP electrodes

Electroencephalography (EEG): EEG and Electromyography (EMG) electrodes were implanted in isoflurane (1.5–3%) anesthetized mice. Carprofen was given post operatively for two days. A prefabricated head mount (Pinnacle Technology Inc., Lawrence, KS) was used to position three stainless-steel epidural screw electrodes. The first electrode (frontal—located over the frontal cortex) was placed 1.5 mm anterior to bregma and 1.5 mm lateral to the central suture, whereas the second two electrodes (interparietal—located over the visual cortex and common reference) were placed 2.5 mm posterior to bregma and 1.5 mm on either side of the central suture. The resulting two leads (frontal–interparietal and interparietal–interparietal) were referenced contralaterally. A fourth screw served as a ground. Electrical continuity between the screw electrode and head mount was aided by silver epoxy. EMG activity was monitored using stainless-steel Teflon-coated wires that were inserted bilaterally into the nuchal muscle. The head mount (integrated 2 × 3 pin socket array) was secured to the skull with dental acrylic. Mice were allowed to recover for at least 14 days before sleep recording.

Local field potential (LFP): LFP, EEG and EMG electrodes were identical to the EEG surgery described above with the following exceptions. A custom-made implant, consisting of two unipolar tungsten electrodes permanently attached to a 2×4 pin socket (Pinnacle Technology Inc., Lawrence, KS), was lowered through a craniotomy with the aid of a stereotaxic apparatus (David Kopff Instruments, Tujunga, CA). The implant was positioned so that the electrodes tips were in the infralimbic (anterior posterior [AP]: +1.9, medial lateral [ML]: -0.4, dorsal ventral [DV]: -3.1; coordinates relative to bregma and midsagittal suture) and prelimbic (AP: +1.9, ML: +0.4, DV: 4.45) cortex. Four epidural, stainless-steel screw-electrodes (Pinnacle Technology Inc., Lawrence, KS) were then positioned on the skull as follows: one recording electrode and LFP reference were placed contralaterally over the frontal cortex, two electrodes over the cerebellum served as an EEG reference and ground. Wire leads attached to each screw were then soldered to output pins on the implant. The implant and leads were covered and secured with dental acrylic.

### Excitatory DREADD

Bilateral injections into the preoptic area (POA) were performed in isoflurane (1.5–3%) anesthetized mice. A 0.5µl microliter syringe needle (Hamilton, Reno, NV) was positioned in the POA, through a craniotomy made in the skull, with the aid of a stereotaxic apparatus (AP: +0.20, ML: ±0.55 DV: -5.65). 200nl of adeno-associated virus 2 (AAV2) containing either control construct (pAAV-hSyn-EGFP; plasmid #50465; Addgene, Watertown, MA) or excitatory DREADD (pAAV-hSyn-hM3D(Gq)-mCherry; plasmid #50474; Addgene, Watertown, MA) was delivered to the POA at 1nl/s using a motorized syringe pump (World Precision Instruments, Sarasota, FL). The syringe needle was left in place for ten minutes before removal. Mice were given 14 days of recovery before EEG implant surgery. Carprofen was given post operatively for two days.

### Clozapine N-Oxide (CNO)

Clozapine N-oxide dihydrochloride (CNO; 2mg/kg; Cat# HB6149; HelloBio, Princeton, NJ) was diluted in in lactated ringers and delivered by intraperitoneal (IP) injections on each day of social defeat, where indicated (**Fig. 2A**), between ZT1 and ZT2 (ZT12=lights off, early light phase). Mice expressing both control construct and excitatory DREADD received CNO. For validation studies (**Fig. 2C**), single IP injections were delivered at ZT 10.

### Sleep Restriction

Sleep restriction was accomplished by one of two methods. In Figure 4, six-hour sleep restriction was conducted by hand and performed by trained observers. Mice were kept awake during the first 6 hs of the light phase (ZT 0–6) once by gentle handling (introduction of novel objects into the cage, tapping on the cage and when necessary delicate touching) and allowed an 18-hr recovery opportunity (ZT 6–0). In Figures 1, 5 and 6 sleep deprivation was accomplished using an automated system (Pinnacle Technology, Inc. Lawrence, KS) which maintained wakefulness by means of a slowly rotating bar in the cage bottom. The bar direction was set to change randomly every 10–20 s. In Figure 2, mice were kept awake for the first 8 hours of the light phase (ZT 0–8) daily for each of the 10-days of social defeat stress. Mice were continuously observed during the sleep restriction by trained observers. In Figure 5 & 6, mice were kept awake for the first 6 hs of the light phase (ZT 0–6) once and allowed an18-hr recovery opportunity (ZT 6–24). Sleep restriction was always carried out in the home cage of the mouse and food and water were available *ad libitum*.

### Social Defeat and Social Avoidance Test

Mice were exposed to three daily, five minute, social defeat sessions (separated by five-minute breaks) for ten days (during the first 2h of the dark period; ZT 12–14). The C57BL/6J mice to be defeated were placed into the home cage of a trained CD-1 mouse (aggressor). Each 5-minute session was with a novel aggressor, no defeated mouse encountered an aggressor more than twice over the 10 days of social defeat stress. Training of aggressor mice was conducted on each of the 3 days prior to testing. Training consisted of the same three, five-minute sessions, but was performed with a C57BL/6J training mouse not used in experiments. Only aggressor mice that displayed at least five incidences of aggression for two consecutive days were used in the study. Social defeat sessions were continuously monitored, and the mice were separated for 10 seconds if excessive aggression (>15 seconds of continuous aggression) was observed. No significant wounding occurred during any of our defeat sessions. Small bite wounds were found on occasion, and they were treated with betadine after the defeat sessions ended. Dim red light (<5 lux) was used for procedures performed during the mouse’s dark period. Novel cage controls were performed with the exact same procedures as social-defeat stress, but with the aggressor mouse moved to a holding cage.

Social avoidance testing was conducted in a 30 × 30 cm arena with a caged (9 × 9 cm), novel, CD1 mouse positioned against the midpoint of one arena wall. Each mouse to be tested was placed in the arena for two consecutive sessions of 3 min. During the first session, the cage was empty; during the second session, the novel CD1 mouse was present. Position in the arena was monitored with video tracking (Noldus Ethovision XT, Leesburg, VA, USA). The arena was cleaned thoroughly between each test. Social interaction ratio, *int*, was calculated as:

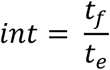

Where *t*_*e*_ = time within 15 cm of the empty cage in the first session, *t*_*f*_ = time within 15 cm of the caged, novel CD1 mouse in the second session; ratio >1.1 = resilient, ratio <0.9= susceptible. Mice with interaction ratios between 0.9 and 1.1 (n=11) were excluded from analysis in the experiments of Fig. 3–6 as follows: Fig. 3, four mice excluded; Fig. 4, four mice excluded; Fig. 5 & 6, three mice excluded.

### EEG/LFP recording/scoring

One week after surgery, mice were moved to an open-top sleep-recording cage and connected to a lightweight tether attached to a low-resistance commutator mounted over the cage (Pinnacle Technologies, Lawrence KS). This enabled complete freedom of movement throughout the cage. Except for the recording tether, conditions in the recording chamber were identical to those in the home cage. Mice were allowed a minimum of seven additional days to acclimate to the tether and recording chamber. Data acquisition was performed on personal computers running Sirenia Acquisition software (Pinnacle Technologies). EEG signals were low-pass filtered with a 30-Hz cutoff and collected continuously at a sampling rate of 400 Hz. LFP signals, and EEG signals collected simultaneously with LFP, were low-pass filtered with a 1000-Hz cutoff and collected continuously at a sampling rate of 2 kHz. In most cases, sleep recordings were conducted in blocks of 8 mice including both treatments and controls. For classification of waveforms, these EEG signals were low-pass filtered offline at 30 Hz. After collection, all waveforms were classified by a trained observer (using both EEG leads and EMG; in 10-sec. epochs) as wake (low-voltage, high-frequency EEG; high-amplitude EMG), NREM sleep (high-voltage, mixed-frequency EEG; low- amplitude EMG) or rapid eye movement (REM) sleep (low-voltage EEG with a predominance of theta activity [6–10 Hz]; very low amplitude EMG). In all studies, individuals performing sleep-stage classification were blind to the experimental conditions and behavioral phenotypes until final analysis. EEG epochs determined to have artifact (interference caused by scratching, movement, eating, or drinking) were excluded from analysis. Artifact comprised less than 5% of all recordings used for analysis.

### Data analysis

Wave incidence analysis has been described previously (*24, 59*). Briefly, analysis was performed using custom written functions in IGOR Pro 8 (WaveMetrics Inc., Lake Oswego, OR). Raw EEG and LFP signals were band pass-filtered in the frequency range indicated using a Butterworth fourth-order band-pass filter (IGOR Pro routine FilterIIR; WaveMetrics Inc., Lake Oswego, OR). Peaks in the filtered data were detected as negative deflections between two zero crossings. The upper 30% of peak amplitudes that occurred in epochs identified as wake were then counted and expressed as peaks per minute (wave incidence). The wave incidence data were binned for graphing and statistical analysis.

Power spectral analysis was accomplished by applying a fast Fourier transform (FFT, 0.1 Hz frequency resolution) to EEG or LFP recordings. Where indicated, spectral power within a frequency band was normalized to 24-h baseline values for each animal or expressed as a percentage of total power (0.5–30 Hz) for each animal. Slow-wave energy, *energy*, was calculated using the 0.5–4Hz (delta) frequency range as follows:

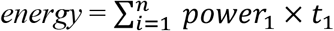

Phase coherence is calculated utilizing the Igor Pro DSPPeriodogram function which can calculate the degree of coherence between the input of two sources, in this case LFP and/or EEG records stored in the same EDF file and on the same time base. According to the Igor Pro literature, the coherence, *coh*, is given by:

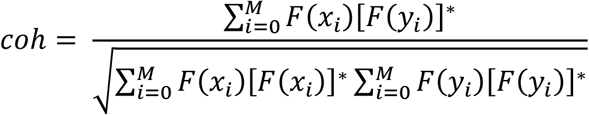

where F(x_*i*_) and F(y_*i*_) are the Fourier transforms of the first and second EEG or LFP data chunks, *i*, for the same time period. The complex conjugate of the Fourier transform is symbolized by the standard notation of [ ]*. The data were recorded at 2 kHz. The data in the coherence plot are analyzed in 10-minute bins made from successive chunks (i) of 2 s each and averaged according to the above equation (*M* = 300 times). Igor Pro also applied a Hanning window to the data to remove edge effects. Frequency data resolution was 0.5 Hz. After Fourier analysis, the real and imaginary parts of each point of coherence on the frequency spectrum, *coh*, were squared and summed to yield a real number values which is the power. This is displayed between 0.5 and 20 Hz (each column on the coherence plot in Fig. 6).

Sleep data were analyzed using one-way or two-way analysis of variance (ANOVA) with repeated-measures when appropriate. Student’s t was used where indicated and for all tests significance was defined as p<0.05. Post hoc analysis was conducted using the Holm-Sidak method which adjusts α to maintain the family-wise error-rate at 0.05. Sample sizes (biological replicates) for each experiment are indicated in the figure legends. An appropriate sample size of 6 was predicted with Type I error rate of 0.05 and Type II error rate of 0.2. Standard deviation and mean difference were estimated as 14.6 and 25 min, respectively, based on previous recordings of C57BL/6J mice obtained in our lab.

### Corticosterone

Cages were ch0anged 24-hours prior to each sample collection at ZT 12 (lights off). All feces from the cage were collected and immediately frozen (-80°C) until assay. Just prior to assay, the fecal sample was homogenized and 50 mg was removed for analysis. Sample preparation and analysis was done using the Cayman Corticosterone ELISA kit (Ann Arbor, Michigan), according to the kit -booklet instructions for fecal extraction and analysis. Each condition was run in duplicate, an average of these values was used in analysis.

### Histology

Under deep isoflurane anesthesia mice were perfused transcardially with 50mL of cold 1M phosphate buffer saline (PBS) followed by 50mL 0.4% paraformaldehyde (PFA). Brains were removed and post-fixed in 0.4% PFA for two days and then transferred to 1M PBS until sectioning. Coronal cryostat sections (25 µm) we transferred to glass slides and air dried. For LFP experiments, sections were stained with cresyl-violet and electrode locations were verified using light microscopy. Brain sections from DREADD experiments were mounted with DAPI-containing mounting medium (nucleic-acid counter-stain; Fluoromount-G, Invitrogen); the presence and location of DREADD expressing cells were verified with laser scanning confocal microscopy (**Fig. S4**; LSM700, Carl Zeiss, White Plains NY). Excitation lasers were 405 nm (DAPI), 488 nm (EGFP), and 561 nm (mCherry).

## Funding

National Institutes of Health grant GM127260 (JCE)

National Institutes of Health grant NS078410 (KNP)

National Institutes of Health grant MD007602

National Institutes of Health grant HL103104 (supported BJB)

National Institutes of Health grant HL007901 (supported EAA)

National Institutes of Health grant HL117929 (supported CLG)

National Institutes of Health grant HL116077 (supported AJB)

### Manuscript Review

Zach Hall and India Nichols-Obande

---

**Fig. S1.**
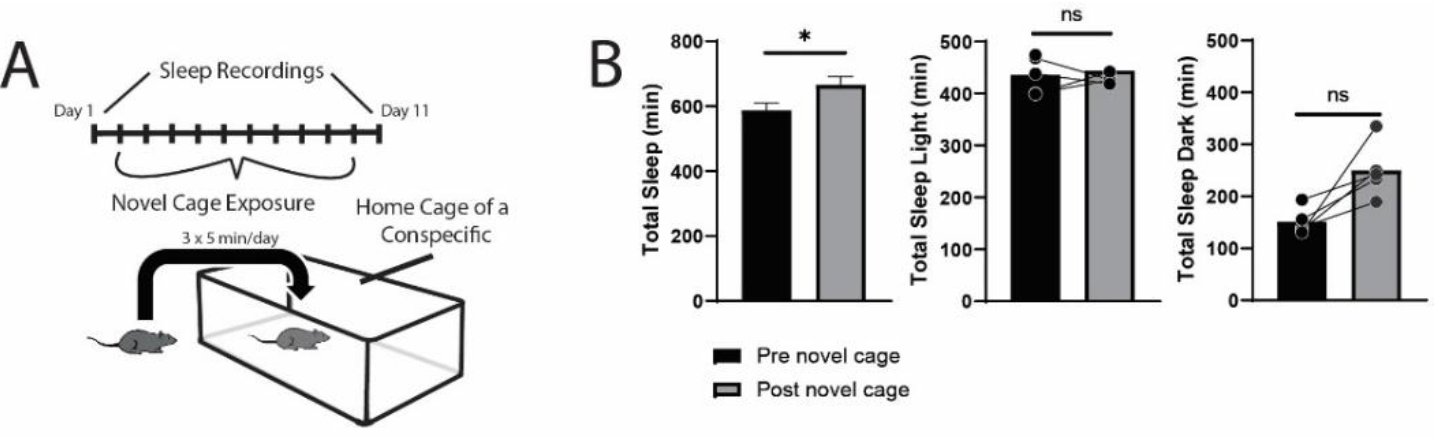
Sleep is not reorganized after exposure to a novel cage. 24-hour sleep recordings were performed both before and following 10-daily exposures to a novel cage (A). This was done to control for the novel environment experienced during social-defeat stress. Total sleep was slightly increased after 10 daily exposures to a novel cage (B) however, no significant differences were detected in the light period or dark period separately. (B) Total sleep— Student’s *t*, t(9) = 2.65, *p* = 0.026; light—Student’s t, *t*(9) = 0.493, *p* = 0.643; dark—Student’s t, *t*(9) = 1.86, *p* = 0.1; n = 10; bars represent mean ± SEM *, *p* ≤ 0.05.

**Fig. S2.**
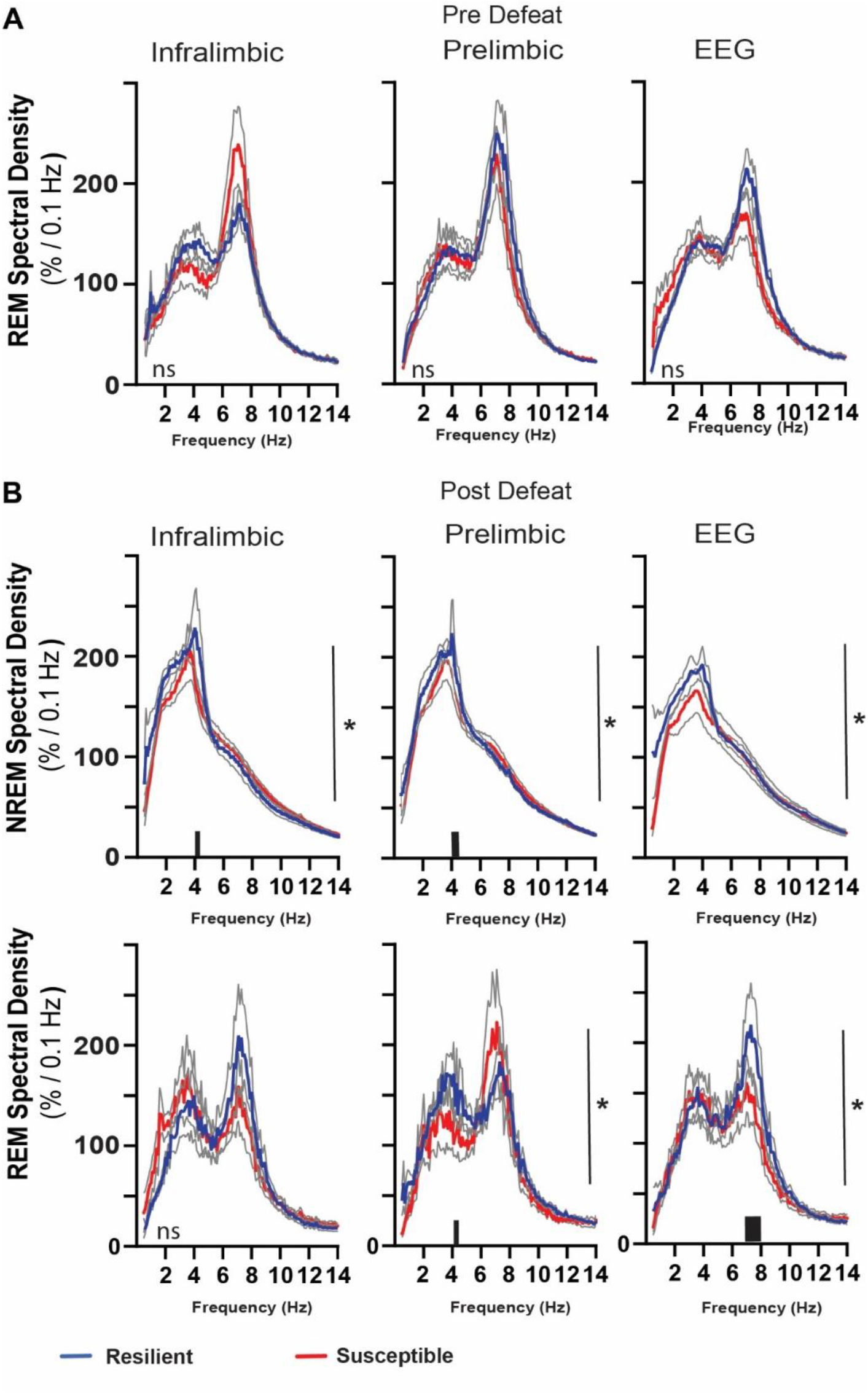
Power density in resilient vs. susceptible animals. Simultaneous local-field potential (LFP)recordings in the ventromedial prefrontal cortex and electroencephalographic (EEG) recordings were obtained from mice before (A) and after (B) ten-days of social defeat stress. REM power density before social defeat stress was not significantly different between resilient (blue) and susceptible (red) mice (A; repeated measures ANOVA). After social defeat stress, resilient animals had increased NREM power density over susceptible animals in frequencies ≤ 4Hz for both LFP and EEG leads. Power in REM power density centered around 4–8Hz also occurred in resilient animals after social defeat stress, compared to susceptible. n = 12. * = significant interaction effect, *p* < 0.05, repeated measures ANOVA. Grey lines indicate SEM; Black bars on x-axis = *p*<0.05, Holms Sidak’s multiple comparison test.

**Fig. S3.**
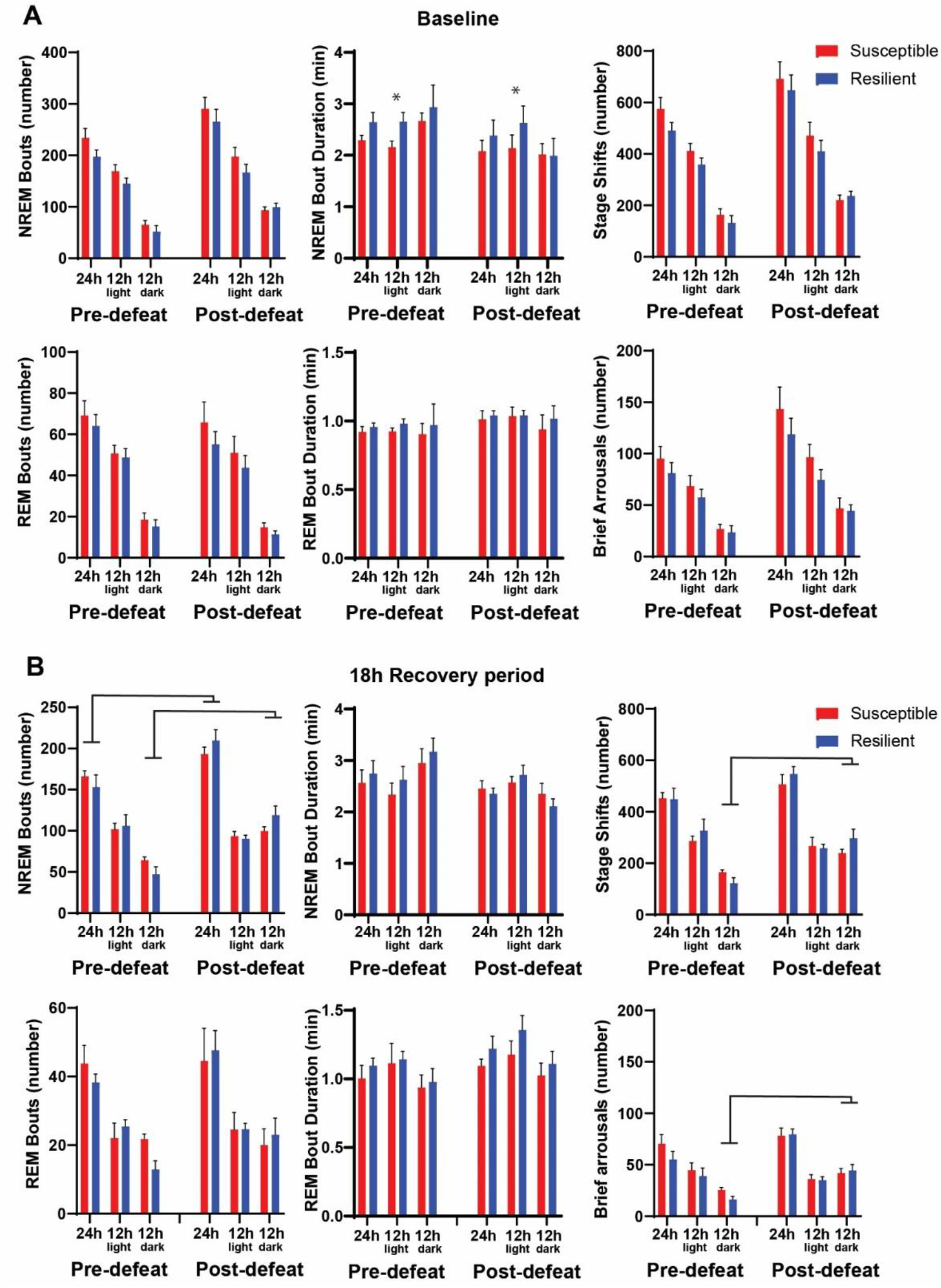
Sleep fragmentation before and after social defeat stress. Sleep fragmentation was assessed from 24-hour recordings during undisturbed sleep and with a six-hour sleep restriction. Recordings were performed both before and after 10-days of social defeat stress (See Fig. 4 A for experimental paradigm). Comparing undisturbed, baseline, sleep recordings revealed slightly longer NREM-bout duration in resilient animals both before and after social defeat stress, but only in the 12-h light period (A, top row). No other differences between resilient and susceptible mice were found during baseline or following 6-hours of sleep restriction (A, B). An overall increase in NREM bout number, stage shifts and brief arousals occurred in both resilient and susceptible animals after social defeat stress. These increases were found when comparing the responses to 6-hours of sleep restriction (B) and were primarily the result of increases during the dark period (B). (A) pre social-defeat stress—*t*(10)=1.87, *p* = 0.046; post social-defeat stress— *t*(10)=2.52, *p* = 0.03; bars represent mean ± SEM; *, *p* ≤ 0.05 Student’s t; n = 13. (B) ANOVA main effect of stress, 24h: bout-number—F(1, 18) = 39.99, *p* < 0.0001; main effect of stress, darks: bout number—F(1, 17) = 14.43, *p* = 0.0047; stage shifts—F(1, 17) = 32.65, *p* < 0.0001; brief arousals—F(1, 7) = 30.82, *p* < 0.0001.

**Fig. S4.**
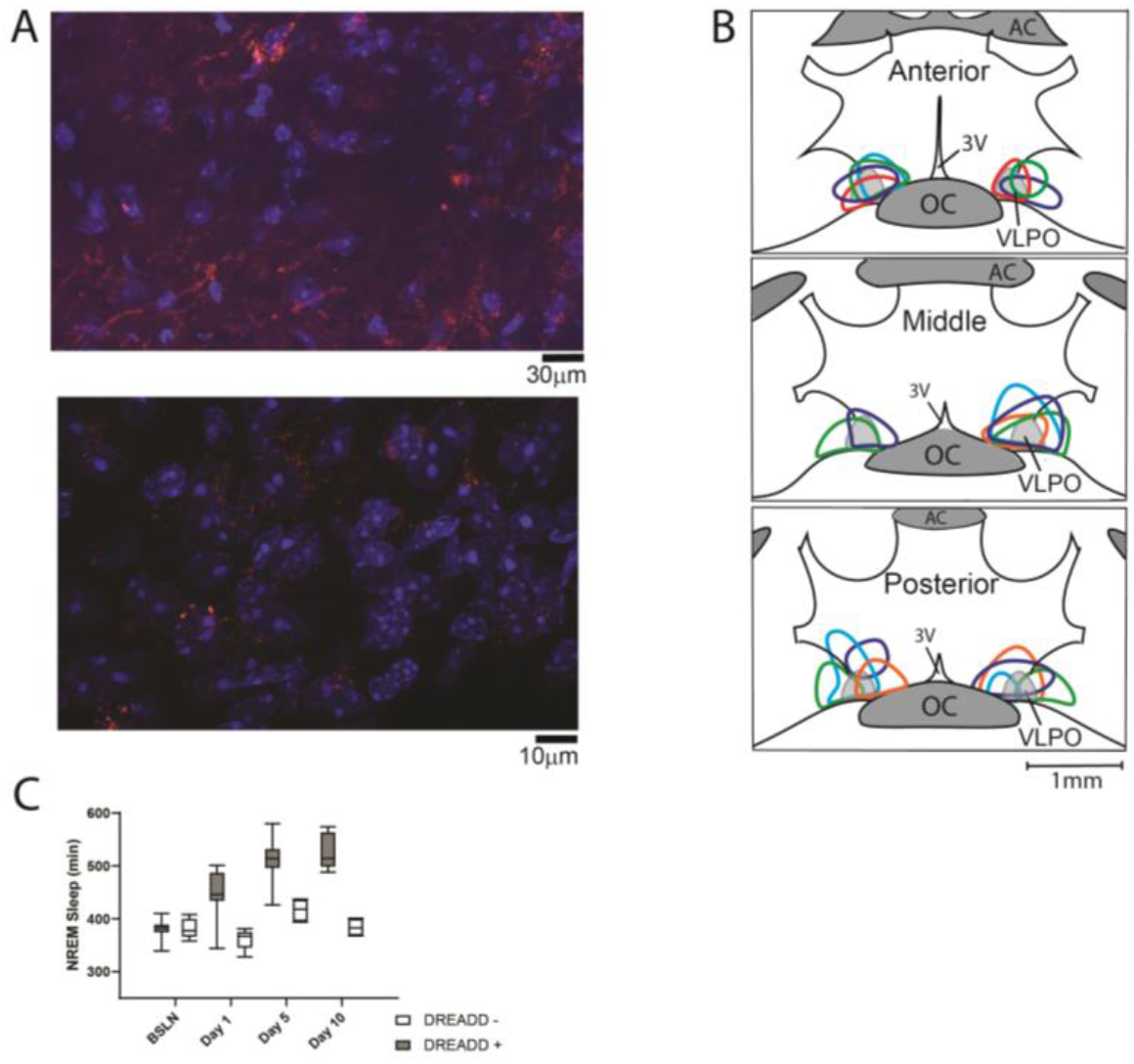
Histological verification of designer-receptor exclusively activated by designer-drugs (DREADD). Verification of pAAV-hSyn-EGFP (control, not shown) or pAAV-hSyn-hM3D(Gq)-mCherry (excitatory DREADD) expression was performed in sections with nucleic-acid counter stain (DAPI). Representative photomicrographs are shown in A (red, mCherry; blue, DAPI). The outer boundary of labeled cell bodies for each mouse included in the study are illustrated in B. Locations are superimposed on representations of hypothalamic coronal sections (∼150 µm spacing; OC, optic chiasm; 3V, third ventricle; AC anterior commissure). Colors are to aid in differentiation and do not identify individual mice. All mice had both increased sleep over the 10 days (C) and labeling that overlapped a portion of the ventrolateral preoptic area (B, VLPO). (C) box and whiskers replot of data in Fig. 2B showing median, 25th to 75th percentiles and min/max values.

## Notes

### Competing Interest Statement

The authors have declared no competing interest.

